# Alpha-band lateralization and microsaccades elicited by exogenous cues do not track attentional orienting

**DOI:** 10.1101/2022.12.12.520080

**Authors:** Elio Balestrieri, René Michel, Niko A. Busch

## Abstract

We explore the world by constantly shifting our focus of attention towards salient stimuli, and then disengaging from them in search of new ones. The alpha rhythm (8-13 Hz) has been suggested as a pivotal neural substrate of these attentional shifts, due to its local synchronization and desynchronization that suppresses irrelevant cortical areas and facilitates relevant areas, a phenomenon called alpha lateralization. Whether alpha lateralization tracks the focus of attention from orienting toward a salient stimulus to disengaging from it is still an open question. In this study, we addressed this question by leveraging the well-established phenomenon of Inhibition of Return (IOR), consisting of an initial facilitation in response times (RTs) for target stimuli appearing at an exogenously cued location, followed by a suppression of that location. Our behavioral data showed a typical IOR effect with both early facilitation and subsequent inhibition. By contrast, alpha was lateralized only in the cued direction, but never re-lateralized in a manner compatible with the behavioral inhibition effect. Importantly, also the initial lateralization towards the cue ocurred too late to account for the behavioral facilitation effect. Furthermore, we analyzed the interaction between alpha lateralization and microsaccades: at the same time when alpha was lateralized towards the cued location, microsaccades were mostly oriented away from the cued location. Crucially, the two phenomena showed a significant positive correlation. These results indicate that alpha lateralization reflects primarily the processing of salient stimuli, challenging the view that alpha lateralization is directly involved in exogenous attentional orienting per se. We discuss the relevance of the present findings for an oculomotor account of alpha lateralization as a modulator of cortical excitability in preparation of a saccade.

## Introduction

The most salient neuronal signal in the human electroencephalogram is the alpha rhythm – an oscillatory pattern over the occipital cortex with a characteristic frequency of around 10 Hz. The distribution of alpha oscillations across the hemispheres is correlated with spatial attention, such that alpha power is typically reduced in the hemisphere contralateral to the attended location relative to the ipsilateral hemisphere. This lateralization is usually induced with cues indicating a task-relevant location. Cue-induced lateralization has been observed in a variety of tasks and paradigms including endogenous cueing (Worden et al., 2000; Busch and VanRullen, 2010; Harris et al., 2018), exogenous cueing (Feng et al., 2017; Keefe and Störmer, 2021), visual working memory (VWM, Sauseng et al., 2009; Boettcher et al., 2021), and visual search (Bacigalupo and Luck, 2019). Collectively, these findings demonstrate that the direction and timing of alpha lateralization coincides with the selection of task-relevant spatial locations. However, recent studies have called into question whether alpha lateralization has a direct causal role in the selection process, or whether it is a secondary consequence of cueing.

According to a long-standing interpretation, cue-induced alpha lateralization reflects the active regulation of cortical inhibition, whereby processing in task-relevant areas (e.g. the occipital cortex contralateral to visual stimulation) is protected from distraction by suppressing task-irrelevant areas (Jensen and Mazaheri, 2010; Foxe and Snyder, 2011). This functional suppression account is based on the well-established inverse relationship between the alpha rhythm and neuronal excitability (Romei et al., 2008), whereby strong alpha power is associated with reduced single-unit firing rates (Chapeton et al., 2019; Haegens et al., 2011), multiunit activity (Bollimunta et al., 2008; Van Kerkoerle et al., 2014), fMRI BOLD signal (Goldman et al., 2002; Mayhew et al., 2013), or contrast perception (Iemi et al., 2017; Balestrieri and Busch, 2022), supporting the idea that alpha oscillations implement spatial selection by decreasing excitability of task-irrelevant cortical areas.

However, a number of studies have found evidence that cannot be easily reconciled with a direct causal involvement of alpha lateralization in cue-related suppression: while alpha lateralization robustly follows the task-relevant location, it is often not affected by the location of irrelevant or distracting stimuli (Noonan et al., 2016; Schroeder et al., 2018; van Moorselaar and Slagter, 2019; Foster and Awh, 2019; Pietrelli et al., 2022). Moreover, Antonov and colleagues (2020) compared the dynamics of alpha lateralization to that of steady state visual evoked potentials (SSVEP), an established neuronal marker of stimulus processing in the early visual cortex. As expected, SSVEPs were amplified at attended locations, but alpha lateralization towards those locations lagged behind the SSVEP effect. Moreover, while alpha power increased in the hemisphere representing the unattended location, this was not associated with SSVEP suppression (Antonov et al., 2020; Gundlach et al., 2020). Collectively, these findings suggest that alpha lateralization might be a secondary consequence of cueing and spatial selection.

Likewise, spatial cueing is often associated with microsaccades – miniature eye movements that occur even when subjects are instructed to keep fixating. Critically, the occurrence and direction of microsaccades is influenced by endogenous (Engbert and Kliegl, 2003) and exogenous cues (Rolfs et al., 2004; Galfano et al., 2004; Lv et al., 2022). Thus, while microsaccades may not be causally necessary for attentional modulation of neuronal responses (Yu et al., 2022), they are modulated as a consequence of attentional selection. Interestingly, recent findings show a strong interdependence between local changes in alpha power and microsaccades (Liu et al., 2022a) or eye movements in general (Popov et al., 2021). By showing such interdependence, these results undermine even more the idea that alpha oscillations are a determinant mechanism of spatial selection. And at the same time, they create the need of better understanding of how alpha oscillations and microsaccades covary throughout, and contribute to, the process of spatial selection.

In the present study, we tested the hypothesis that alpha lateralization is a consequence of cue processing that is not directly involved in attentional orienting. Most previous studies have been designed to induce attentional orienting at the cued location, making it impossible to distinguish cue processing and orienting. Here, we leveraged the inhibition of return (IOR) phenomenon, where attention is oriented away from an exogenously cued location (Posner et al., 1985; Klein, 2000). In a typical Posner task, in which an exogenous, uninformative cue is followed by a target after a variable delay, reaction times show a robust two-stage pattern: short-lived facilitation at short delays with faster responses to targets at the cued position followed by long-lasting inhibition at delays longer than ∼ 220 ms with faster responses at the uncued position (Samuel and Kat, 2003). This pattern is believed to reflect the time course of attentional deployment, with an initial orientation towards the cued position, and subsequent disengagement and re-orientation towards the uncued position (Klein, 2000; Klein and Hilchey, 2011). Accordingly, if alpha lateralization reflects spatial orienting, the time course and direction of alpha lateralization should mirror the behavioral effect of IOR, by showing an initial facilitation for the cued location (increase in alpha power ipsilaterally, decrease contralaterally), followed by a pattern reversal corresponding to the inhibition of the cued position. Alternatively, if alpha lateralization were only a secondary consequence of attentional orienting associated with cue processing, the direction of lateralization should only reflect the cued location. Moreover, given that the typical exogenous Posner cueing task is accompanied by microsaccades away from cued position at approximately the onset of the IOR (Rolfs et al. 2004, Galfano et al. 2004, Lv et al. 2022), we also aimed to characterize the interaction between cue-induced shifts in alpha lateralization and microsaccades.

## Methods

### Pre-registration and Participants

The present study was conducted and analyzed as described in the pre-registration (aspre-dicted.org), unless stated otherwise. We had planned to collect data from 20 participants after exclusion based on: 1) poor behavioral performance (False Alarm Rate, FAR > .20); 2) incomplete sessions; 3) poor EEG data quality (n trials rejected for artifact > 15%). We recorded data from 23 participants. Data from three participants were excluded due to high FAR and data from one participant were discarded due to poor eye-tracking data quality, a criterion that had not been pre-registered. The final sample was composed of 19 participants (F=15, M=4, mean age=22.5 yrs, age range 19-28). All participants gave written consent for their participation, reported normal or corrected to normal vision, and were compensated either with money (8 EUR/hour) or with course credits. The study was approved by the ethics committee of the Faculty of Psychology and Sports Science, University of Münster.

### Apparatus

Recordings took place in a dimly-lit, sound-proof cabin. Participants placed their heads on a chin-rest and could adjust the height of the table to be seated comfortably. Stimuli were generated using MATLAB 2019a (mathworks.com) and Psychtoolbox (Brainard, 1997; Kleiner et al., 2007). The experiment was controlled via a computer running Xubuntu 16.04, equipped with an Intel Core i5–3330 CPU, a 2 GB Nvidia GeForce GTX 760 GPU, and 8 GB RAM. The experiment was displayed on a 24” Viewpixx/EEG LCD Monitor with 120 Hz refresh rate, 1 ms pixel response time, 95% luminance uniformity, and 1920*1080 pixels resolution (vpixx.com). Distance between participant eyes and the monitor was approximately 86 cm. Eye-movements were monitored using a desktop-mounted Eyelink 1000+ infrared based eye-tracking system (sr–research.com) set to 500 Hz sampling rate (binocular). EEG was recorded with a Biosemi Active Two EEG system with 65 Ag/AgCl electrodes (biosemi.nl), set to 1024 Hz sampling rate. Sixty-four electrodes were arranged in a custom-made montage with equidistant placement (“Easycap M34”; easycap.de), which extended to more inferior areas over the occipital lobe than the conventional 10–20 system (Oostenveld and Praamstra, 2001). An additional external electrode was placed below the left eye.

### Stimuli and Experimental Procedure

For an overview of the stimulus arrangement see Figure 1A. All stimuli were presented on a medium gray background (52.2 cd/m2). Two placeholders indicating target locations were positioned at 8 dva to the left and right of the central fixation marker (diameter = 1 dva, black and white, 0.2 cd/m2 and 102.3 cd/m2; see Thaler et al., 2013). The cue was an amplification of one of the two target location placeholders, i.e. thicker and darker (16.6 cd/m2, linewidth = 0.08°). The target was a dark square (diameter = 0.1°, 23.7 cd/m2) centered in one of the two target locations. After the target, only the placeholders and the fixation cross were presented until the response. A small colored rectangle (side = 0.2 dva) presented in the center of the screen served as feedback at the end of a trial (yellow for correct, blue for incorrect responses).

**Figure 1:**
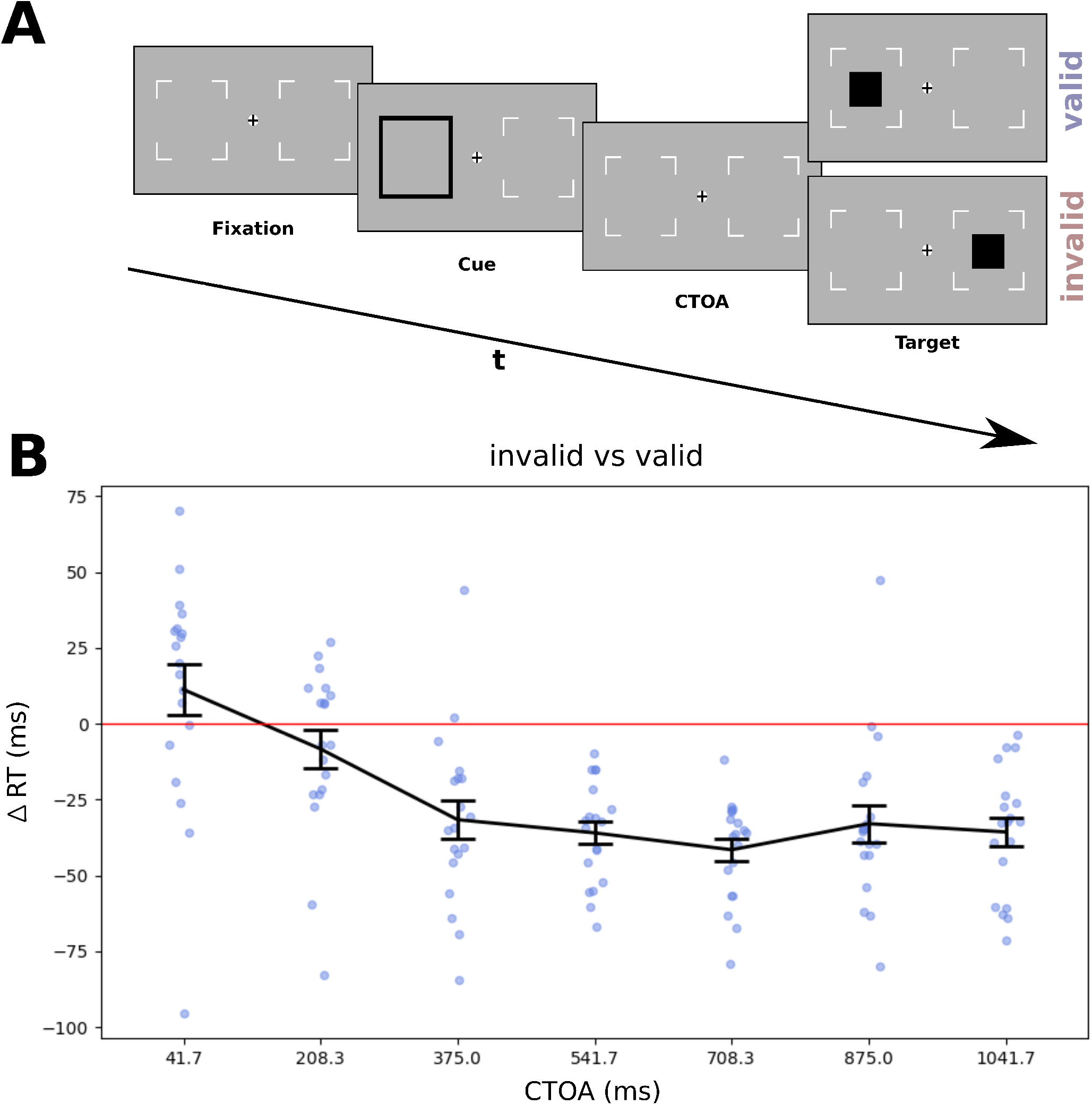
**A)** Illustration of stimuli and procedure. Every trial started with a fixation cross flanked by 2 placeholders. After a jittered time interval, an exogenous cue (the black contour) flashed for 100 ms at one of the two positions. After a variable cue-target onset asynchrony (CTOA) a target (a black square) appeared for 33 ms either at the cued (valid) or uncued (invalid) position, and participants had to detect it by a speeded button press on a gamepad. **B)** RTs difference between invalid and valid cues as a function of CTOA; positive values indicate facilitation, negative values indicate inhibition of the cued position.. Error bars represent SEM..

The study comprised a single recording session of approximately 1.5 h-duration to collect a total of 980 trials. Prior to the experiment, ten to thirty practice trials were presented. Each session was divided into 7 blocks of 140 trials, separated by breaks (self-paced, but at least 30 s), with each block containing a small break (self-paced, but at least 10 s) after 70 trials. The time course of IOR was sampled through 7 cue-target onset asynchronies (CTOAs) ranging from 42 ms to 1050 ms in steps of 168 ms with 28 trials per CTOA and validity condition (784 trials in total). Additionally, 196 catch trials (20%) without a target (from hereafter “no target” condition) were presented, equally distributed across blocks. Cue position, target position and CTOAs were counterbalanced within each block.

The trial sequence is illustrated in Figure 1A. Each trial started with a fixation cross and two placeholders for 800 to 1200 ms (randomized across trials), followed by a non-informative visual cue (cue validity: 50%) flashed for 100 ms. Except for no-target trials, the target was presented after a variable CTOA for 33 ms either at the cued location (valid) or uncued location (invalid). Both placeholders and the fixation marker remained on-screen until the participant’s response. Participants were asked to press a button as fast and accurately as possible within 2 seconds whenever they detect a target regardless of its location. After response, participants received feedback indicating whether their response was correct. Subsequently, a blank screen was presented as inter-trial-interval for a random duration from 1000 to 1500 ms.

Correct fixation was monitored online, and trials in which the gaze drifted away from the central fixation marker for more than 2 dva were interrupted and repeated at the end of the block. The same procedure was applied in trials containing button presses before the target appeared on screen. A second fixation assessment was performed offline, leading to the exclusion of an average of 0.2% (SD=0.3) of trials in which the online fixation control was inaccurate.

### Behavioral analysis

Data preprocessing was implemented in R 4.1.3. For each participant, RTs below 200 ms and above 4 SD were excluded from further analyses. RTs were z-scored based on the mean and standard deviation across all trials, and the time course of IOR was computed by averaging RTs separately for every CTOA and validity condition. Statistical analyses were implemented in python 3.8 (python.org) and pingouin (pingouin-stats.org), using a repeated-measures ANOVA with the within-factors validity (valid, invalid) and CTOA.

Plots representing the evolution of RTs across CTOAs show the difference between invalid and valid trials, so that positive values indicate facilitation (shorter RTs) for the cued position and negative values indicate inhibition (longer RTs).

### EEG preprocessing

EEG preprocessing was performed with EEGLAB (Delorme and Makeig, 2004) and custom scripts running on MATLAB 2021a (mathworks.com). Data were first downsampled to 256 Hz and re-referenced to common average. The continuous data were then high-pass filtered at .8 Hz and low-pass filtered at 40 Hz, and then epoched from -1000 to 2000 ms time-locked to cue onset. A first set of trials were rejected based on extreme amplitudes (+/-500 *µV*) or on joint probability (function *pop_jointprob* in EEGLAB, local threshold 9, global threshold 5). ICA was then performed, and the IClabel algorithm (Pion-Tonachini et al., 2019) was used to classify components in the categories “Brain”, “Muscle”, “Eye”, “Heart”, “Line Noise”, “Channel Noise” and “Other” based on their spatial topography. Components belonging to non-brain sources with a probability above .5, as well as components labeled as “Brain” with a probability below .1 were excluded. After the artifactual components had been removed, noisy channels (whose SD>3.5) were spherically interpolated. Due to good data quality after preprocessing, only one channel in one subject needed interpolation. Finally, remaining trials including amplitudes larger than +/-150 µV were rejected (on average 2.6% of trials per subject; SD=3.4).

### EEG data analysis

The analysis aimed to investigate the dynamics of exogenous orienting and re-orienting indicated by lateralized alpha oscillations. Given that both cues and targets were themselves lateralized stimuli, and giventhat stimulus-evoked potentials comprise substantial power at alpha frequencies (Schürmann and Başar, 2001), it was necessary to remove these stimulus-evoked responses from the signal. To this end, the average across trials (i.e. the evoked potential) was subtracted from each single trial prior to the time-frequency analysis, separately for each cue position x target position x CTOA condition. Thereby, this analysis ensures that the remaining alpha lateralization is not confounded by the cue-evoked or target-evoked sensory response (see Klimesch et al., 1998). As a result, no alpha-band lateralization is visible following target onset in Figure 2, underlying the success of this procedure.

**Figure 2:**
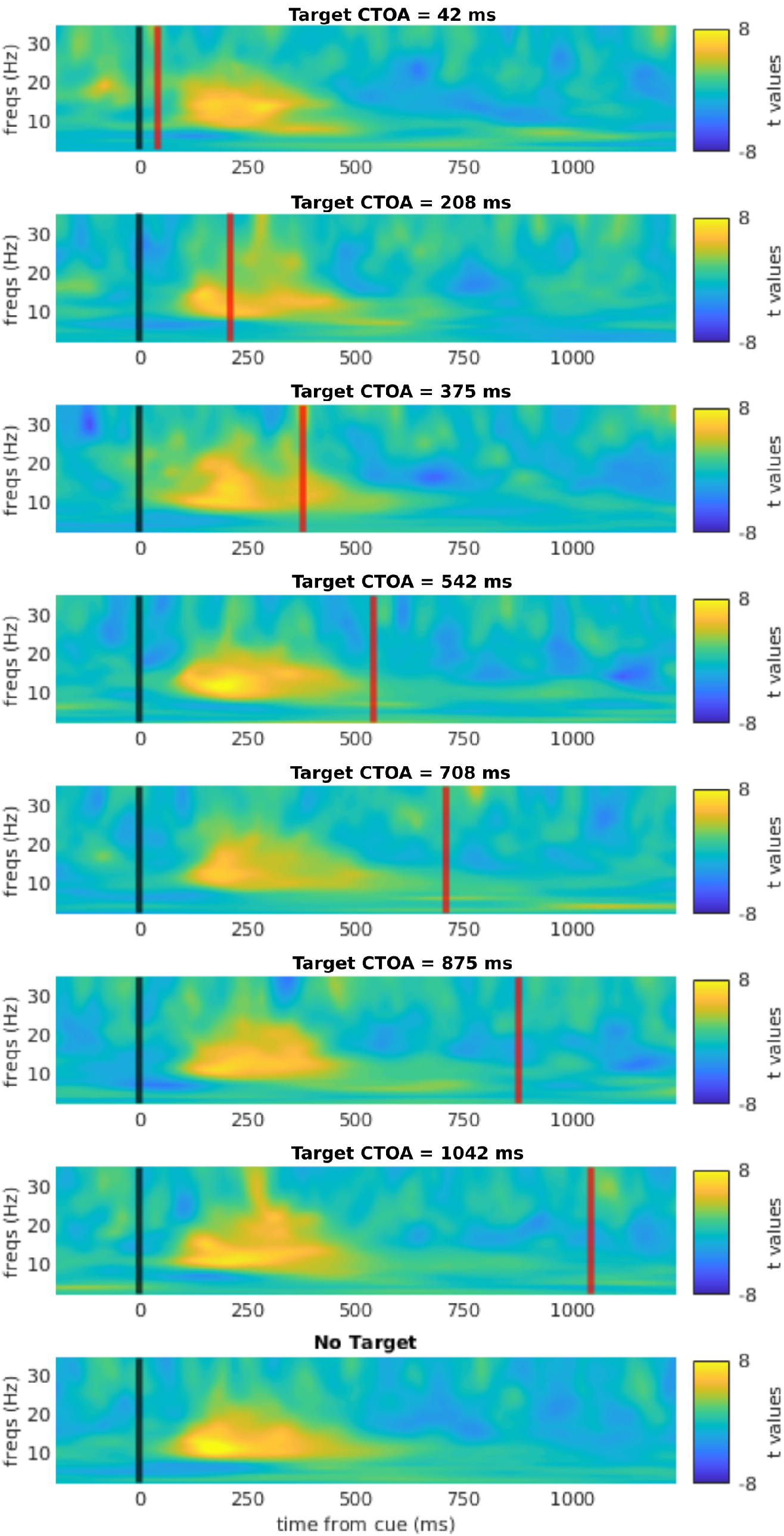
Time-frequency representation of lateralized power in response to the exogenous cue, collapsed across validity conditions. Note that the averaged ERP had been subtracted from the data before time frequency analysis, thereby removing the signals evoked by cue and target onsets. In all subplots, the black line at 0 indicates cue onset, whereas the red line indicates the target onset (except for no-target trials).

The time frequency analysis was performed using a wavelet transform with 34 wavelets over a frequency range between 2 and 35 Hz, full-width at half-maximum (FWHM) = 2/f (Cohen, 2019).

Lateralized power was computed as follows:

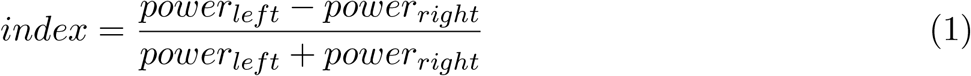

Where *power* is the spectral power (averaged across trials) for each frequency, in the left or right ROI. The ROIs for computing lateralization were two homologous subsets of five left and right occipital channels already shown to maximize alpha lateralization in a previous study (Balestrieri and Busch, 2022). Cue-induced lateralization was then computed as the difference in lateralization index between left and right cued trials. Thus, conventional cue-induced alpha lateralization would take positive values, indicating stronger power ipsilateral vs contralateral to the cue.

The bulk of the EEG analysis focused on the no-target condition, for which the entire post-cue interval was guaranteed to be free of any target-evoked signals. Alpha-band lateralization was further quantified by averaging across frequencies from 8 to 13 Hz. Cue-evoked lateralization was tested for significance with t-tests against 0, under the null hypothesis of lack of any lateralization. We corrected for multiple comparisons through cluster permutation tests (see *cluster permutations tests*).

### Microsaccades

Analyses of microsaccades had not been planned a priori, and were hence not included in the original pre-registration. As for alpha lateralization, only no-target trials were analyzed to avoid target-related confounds. The eye-tracking data in X-Y screen coordinates from both eyes were averaged and microsaccades were detected using the function *pop_detecteyemovements* from the EYE-EEG toolbox (Dimigen et al., 2011) using the microsaccade detection algorithm by Engbert and Mergenthaler (2006). Lambda was set to 5, and the minimal microsaccade duration was set to 3 samples (11 ms). Raw data were smoothed to suppress noise. Furthermore, all eye movements exceeding 1 dva were excluded. Microsaccades in the direction of the cued location were labeled as “towards” and microsaccades in the opposite direction were labeled as “away”. Microsaccadic rate was computed with a linear moving average of 200 ms. As a final step, we computed a metric of microsaccadic rate bias (from hereafter “MS-bias”), as the difference between the rate of microsaccades towards and away from the cue.

### Mutual relationship between alpha lateralization, RTs and MS-bias

We reasoned that if there was any connection between alpha lateralization, RTs and MS-bias, they should co-vary across time. Hence we computed time point by time point Pearson correlations between these variables based on the no-target condition. Alpha lateralization and MS-bias were only quantified for no-target trials. Furthermore, alpha lateralization and MS-bias were subsampled to adjust for the sparse sampling of RTs (i.e. with just 7 CTOAs) as follows.

Alpha lateralization was averaged across 100 ms long time windows, each starting 80 ms after each corresponding CTOA. We reasoned that target processing would be most affected by the magnitude of alpha lateralization at the approximate latency of target-processing in early visual cortex (Iemi et al., 2019), as opposed to target-appearance on the screen. MS-bias was averaged across 100 ms long time windows immediately preceding each corresponding CTOA. Here, we reasoned that target processing would be most affected by the presence of microsaccades (and their direction) immediately before target onset. Pairwise correlations between alpha lateralization, MS-bias, and RTs were computed for all latency-pairs to analyze for each signal at a given CTOA how and when this signal was associated with any of the other signals (e.g. whether alpha lateralization at the earliest CTOS was associated with RTs at later CTOAs). We also computed a sample-by-sample correlation between alpha lateralization and MS-bias using the original data. In all the aforementioned analyses, results were corrected for multiple comparisons via cluster permutation.

### Cluster permutation tests

Cluster permutation tests were used to correct for multiple comparisons. For the tests involving correlations between different time series, we used the procedure detailed by Kluger et al. (2021). The procedure was analogous for (1) correlations between RTs and MS-bias rate, (2) RTs and alpha lateralization, and (3) alpha lateralization and MS-bias. For each timepoint, we correlated the vector of inter-subject values belonging to the first time series (e.g. alpha lateralization) with the paired vector of the other time series (e.g. MS-bias). This yielded a set of Pearson *r* and corresponding p values. In order to correct for multiple comparisons, we applied a cluster permutation approach as follows. We first detected the significant clusters in our data and obtained the cluster masses by summing together the *r* values of the contiguous significant points (p < .05). Then we reiteratively shuffled (N=1000) the participant order in both the time series under test. For each iteration, we stored the absolute highest cluster mass value, yielding a surrogate distribution of cluster masses. We rejected H0 for each cluster mass in the real data that was below the 2.5^th^ or exceeded the 97.5^th^ percentile (*α* = .05, two-tailed) of the surrogate distribution of cluster masses.

For the test against 0 used for assessing lateralization, or MS-bias, we also implemented cluster permutation tests ad hoc. T-tests against 0 were performed for the whole time course. We defined clusters by selecting those t-values contiguous in time and exceeding the critical threshold (*α* = .05) on both tails. From each cluster we obtained a cluster statistic by summing together all the t-values belonging to the cluster, i.e. characterized by contiguity in space (electrodes) and time. To correct for multiple comparisons, for 1000 repetitions, we multiplied a random participants’ subset by -1, and repeated the aforementioned steps in order to obtain a distribution of cluster statistics under the null hypothesis. We rejected the null hypothesis if a cluster statistic was below the 2.5^th^ or above the 97.5^th^ percentiles of the permutations.

## Results

### Behavior

To confirm the existence of an IOR effect, we subjected the RTs to a repeated-measures ANOVA with the within-factors validity (valid, invalid) and CTOA. The RT difference between invalid and valid conditions as a function of CTOA is shown in Figure 1B.

The pattern shows the typical initial facilitation and subsequent inhibition at the cued location, and was confirmed statistically by the ANOVA, which showed, on top of the main effect of validity (F_(1, 18)_=36.103, p<.001, *η*^2^=.667) and of CTOA (F_(6,108)_=26.634, p<.001, *η*^2^=.597) the expected strong validity x CTOA interaction effect (F_(6,108)_=17.431, p<.001, *η*^2^=.492). For each CTOA, we tested the difference between invalid and valid conditions via one-tailed paired samples t-tests. The choice on one-tailed tests was justified by the specific directionality expected: shorter RTs for the validly cued position at the first CTOA, and longer RTs for all the others. The first CTOA showed a trend toward significance (t_(18)_=1.709, p_(uncorrected)_=.052), whereas all the other CTOAs after 225 ms (i.e. the typically observed onset of IOR) showed a strong advantage for the uncued position (all t<-5.729, all p<.001, Bonferroni corrected). This pattern of results shows that we successfully elicited IOR in our participants.

### Spectral and temporal pattern of cue-induced lateralization

We characterized the temporal and spectral profile of lateralized responses to the spatial cues, as presented in Figure 2. For each CTOA and for the no-target trials, we computed the lateralization index for the whole time window of possible target appearance (up to 1050 ms post cue). In all conditions, cue-induced lateralization “towards” the cued location ramped up at around 100 ms after cue onset and leveled off at around 600 ms. This response was most prominent in the alpha band, but extended to higher frequencies in the beta band. This pattern was observed consistently for all CTOAs and both validity conditions, confirming that our exogenous cueing paradigm successfully elicited a relative increase in power at ipsilateral channels relative to the cue. However, we did not observe any re-lateralization “away” from the cue at longer latencies that would mirror the behavioral inhibition of return effect at longer CTOAs. This mismatch between patterns of behavioral data and lateralized power was investigated more systematically in the subsequent analyses.

### Time course of alpha lateralization and its relation to RTs

Given that our a priori hypothesis targeted the alpha band, which was also the band with the strongest lateralized effect in the current analysis, we focused on the power averaged in the 8-13 Hz range, and restricted the analysis to no-target trials, which were unconfounded by any target-evoked signals. The resulting time course showed pronounced and sustained alpha lateralization towards the cue position (i.e., increase ipsilateral and decrease contralateral to cue) starting at cue onset, peaking at ∼250 ms and fading away after 650 ms (cluster p<.001, Figure 3B). Importantly, lateralization did not invert *away* from the cue position at longer latencies.

**Figure 3:**
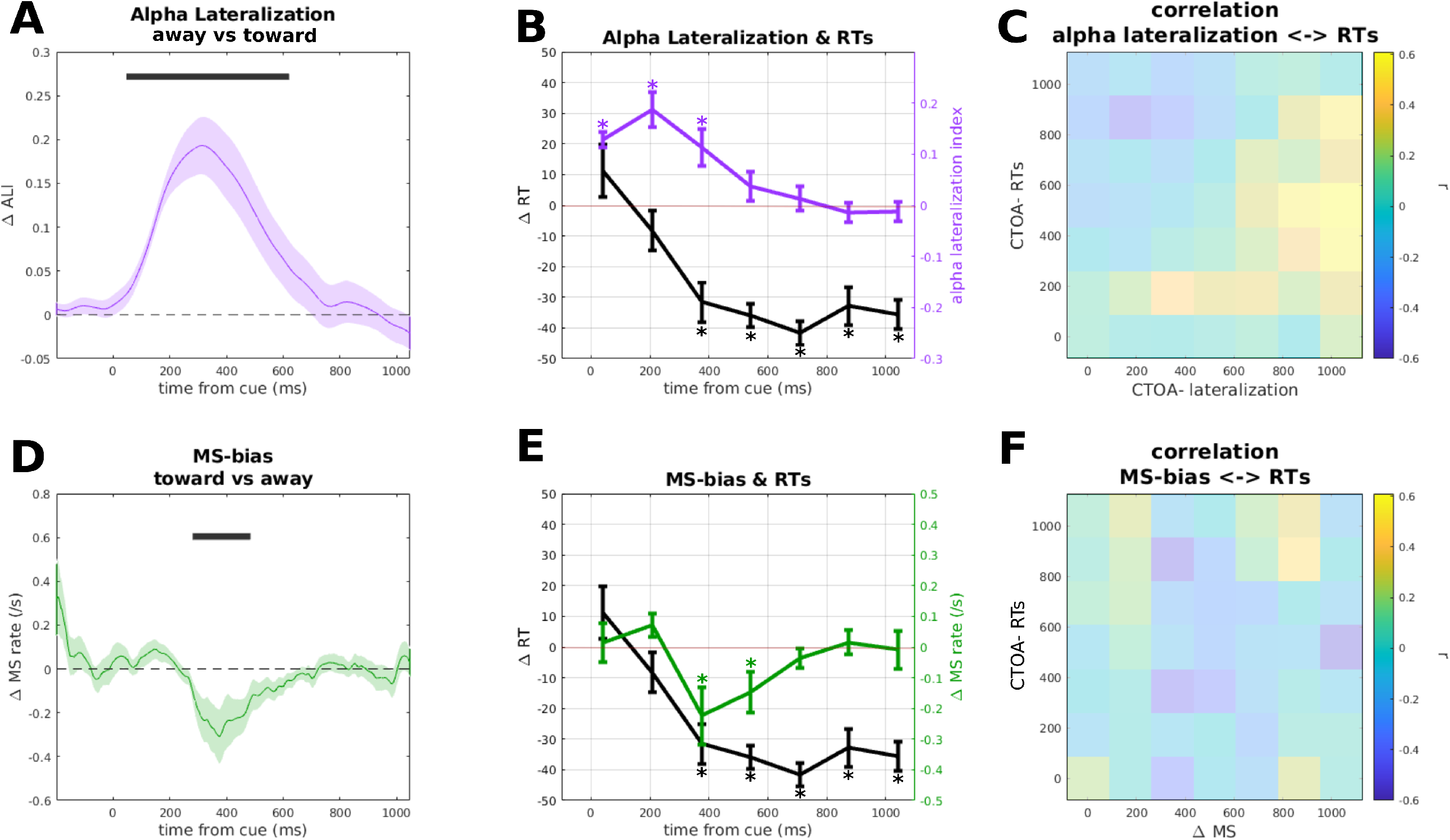
**A)** Lateralization in the alpha band (8-13 Hz). Positive sign indicates increase in alpha ipsilaterally and decrease contralaterally. The dark gray bar indicates the significant cluster tested against 0. Shaded area represents SEM. **B)** Comparison between RTs and alpha lateralization (downsampled to match the RT time course (see Methods). The different color-coded y axes are set to different scales to allow a direct comparison between alpha lateralization and RTs. Error bars represent SEM. Asterisks indicate significant t-tests against 0. **C)** Inter-subjects correlation matrix between alpha lateralization and RTs. The transparency masking indicates that no test survived correction for multiple comparisons. **D)** Microsaccade (MS)-bias, towards (positive) and away (negative) from the cued location. The dark gray bar indicates the significant cluster tested against 0. Shaded area represents SEM. **E)**: comparison between RTs and MS-bias(downsampled to match the RT time course) (see Methods). The different color-coded y axes are set to different scales to allow a direct comparison between MS-bias and RTs. Error bars represent SEM. Asterisks indicate significant t-tests against 0. **F)** Inter-subjects correlation matrix between MS-bias difference and RTs. The transparency masking indicates that no test survived correction for multiple comparisons.

**Figure 4:**
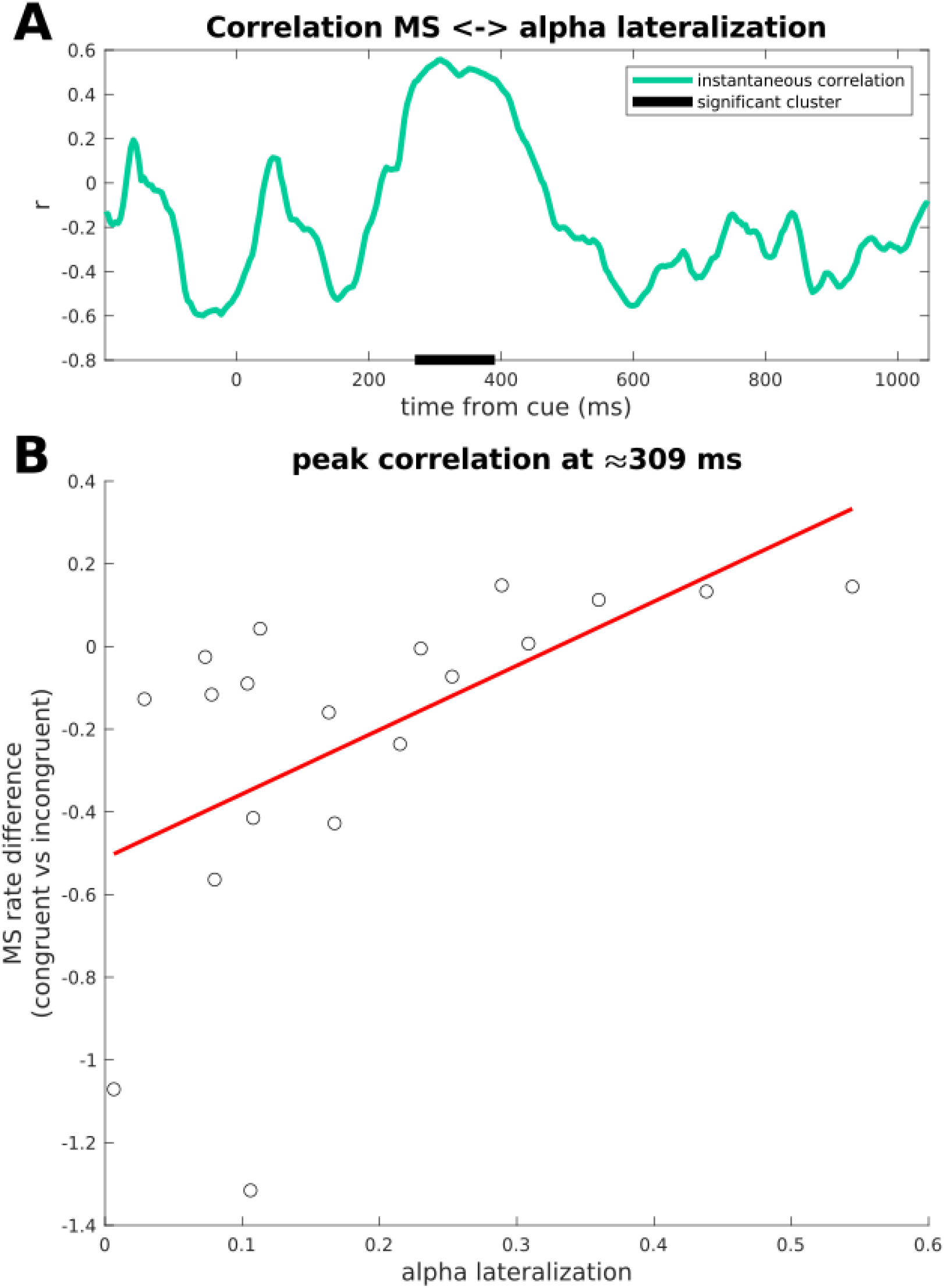
**A)** Correlation over time between alpha lateralization and MS-bias. The dark gray bar indicates the significant cluster after cluster permutation test. **B)** Peak of the positive correlation at ∼ 309 ms.

To compare the pattern of alpha lateralization across time to the pattern of response times across CTOAs, we averaged lateralization in no-target trials within seven time bins coinciding with the seven CTOAs in target trials. Thus, this sequence of lateralization averages in no-target trials reflects the momentary lateralization at the latencies when targets were presented in the different CTOA conditions, allowing for a direct comparison with the sequence of RTs at the corresponding CTOAs (Figure 3B). While the RT time course showed behavioral facilitation at the cued location only at the earliest SOA (42 ms), alpha lateralization towards the cue continued to increase and peaked as late as 200 ms. T-tests against zero performed on each CTOA for alpha lateralization confirm the pattern of results described beforehand, with the first CTOAs showing values significantly higher than 0 (CTOA=42, t_(18)_=8.576, p<.001; CTOA=208, t_(18)_=5.468, p<.001). Furthermore, while the RT time course showed a clear IOR effect at all but the earliest CTOA, alpha continued to be lateralized towards the cue until 375 ms (t_(18)_=3.181, p=.005) and then returned to baseline (CTOA=542, t_(18)_=1.271, p=.219). Thus, while the inhibition of return was in full swing, alpha lateralization did not show a corresponding reversal away from the cue, further confirmed by the last 2 CTOAs which did not diverge significantly from 0 (CTOA=875, t_(18)_=-.718, p=.482; CTOA=1042, t_(18)_=-.677, p=.507).

We reasoned that the apparent mismatch between the time courses of RTs and alpha lateralization might have been influenced by inter-individual variability in either variable. To address that, we conducted a correlation analysis to test if individuals with stronger behavioral IOR showed stronger lateralization in either direction. These correlations were computed between behavioral IOR effects at all CTOAs and lateralization at all latencies, resulting in a 7 × 7 correlation matrix. If stronger lateralization away from the cued location was associated with stronger behavioral IOR at the same time, this would show as a positive correlation in Figure 3C. If stronger lateralization was associated with IOR at a later time, this would show as a correlation above the diagonal. This analysis did not show any significant correlations along the diagonal (Figure 3C). Moreover, although a few correlations were significant (for example positive correlations below the diagonal), none of them survived correction for multiple comparisons.

To sum up, alpha power was strongly lateralized towards the exogenous cue, with no evidence for a later re-lateralization away from the cue mirroring the strong behavioral inhibition of return effect.

### Time course of microsaccades and its relation to RTs

We also examined the effect of exogenous cueing on microsaccadic rate bias (MS-bias), computed as the difference between microsaccadic rate towards and away from the cued location. Again, this was based on the no-target condition. As for the alpha lateralization, we first tested MS-bias against 0 for the whole time window following the cue, revealing a directional bias away from the cued location (Figure 3D), starting around 250 ms, peaking at ∼370 ms and fading away around 600 ms (cluster p=.014). While the onset of this directional bias is compatible with the onset of IOR, MS-bias then fades away after 600 ms, whereas the IOR continues until after 1000 ms.

To compare MS-bias to the pattern of response times across CTOAs, we again averaged MS-bias in no-target trials within seven time bins coinciding with (i.e. just preceding) the seven CTOAs in target trials (Figure 3E). This direct comparison shows an initial trend toward significance for MS-bias congruent to the cued location (CTOA=208, t_(18)_=1.812, p=.087), followed the MS-bias away from the cue at ∼400 ms, already shown by the cluster permutation analysis (CTOA=375, t_(18)_=-2.39, p=.028; CTOA=544, t_(18)_=-2.239, p=.038), to return to baseline in the last two CTOAs (CTOA=875, t_(18)_=.377, p=.711; CTOA=1042, t_(18)_=-.153, p=.883). The correlation between RTs and MS-bias (Figure 3F) did not yield significant results.

### Correlation between MS-bias and alpha lateralization

At first glance, it seems that alpha lateralization and MS-bias showed diametrically opposite directions: the pattern of microsaccades showed a bias away from the cued position, whereas alpha lateralization was directed towards the cued location, although these patterns show a remarkable temporal overlap. Accordingly, a point by point correlation across subjects between the two time courses yielded a cluster of positive correlation with onset at ∼270 ms and offset at ∼390 ms (cluster p =.008), peaking at ∼309 ms (r=.557; p=.013), indicating that subjects with stronger lateralization towards the cue had less MS-bias away from the cue.

## Discussion

In the current study, we tested whether alpha lateralization tracks exogenous spatial attention, and specifically inhibition of return (IOR) in a typical exogenous Posner cueing task. For this purpose, we compared the time course of alpha lateralization after cue onset with the behavioral time course of IOR across seven cue-target onset asynchronies (CTOAs).

As expected, we found the typical IOR effect in behavior (Klein, 2000; Samuel and Kat, 2003) with facilitation at the cued position at short CTOA followed by strong and long-lasting inhibition of the cued position at long CTOAs (Fig. 1B). The IOR is probably not a unitary phenomenon: depending on the specifics of the task and paradigm, it may result from input-based (i.e. sensory, attentional), or output based (oculo-motor) mechanisms (Berlucchi, 2006; Dukewich and Klein, 2015). The experimental paradigm used in the present study, specifically the use of manual responses and tonic suppression of saccades, has been demonstrated to elicit predominantly input-based IOR effects (Klein, 2000; Taylor and Klein, 2000; Klein and Hilchey, 2011; Redden et al., 2021). For instance, several studies have demonstrated that, under these conditions, IOR affects not only response times but also visual sensitivity (i.e. d’ Handy et al., 1999; Ivanoff and Klein, 2006). Furthermore, (McDonald et al., 2009) investigated the N2pc (an event-related potential indicating a shift of attention) in an adaptation of the classical IOR paradigm, and found that the N2pc was reduced for targets presented at recently attended locations. When no target appeared, the N2pc was even directed away from the previously attended location, thus confirming an attentional mechanism. Thus, we conclude that the behavioral IOR effect found in the present study reflects initial attentional orienting towards the cued position, followed by re-orienting of attention away from and inhibition of the cued location.

Given the hypothesized role of alpha lateralization in modulation of input gating under endogenous (Foxe and Snyder, 2011) and exogenous attention (Feng et al., 2017; Keefe and Störmer, 2021), we expected that alpha lateralization would show a time course and direction consistent with the time course of behavioral effects: an increase in alpha ipsilaterally and a decrease contralaterally to the cue at early latencies (<225 ms after cue onset), followed by a reversed lateralization pattern at longer latencies. Indeed, we observed an initial, strong alpha lateralization towards the cue, in agreement with previous studies (Keefe and Störmer, 2021; Feng et al., 2017, ; Fig 2). Nonetheless, this lateralization emerged only after behavioral facilitation had already disappeared (Fig. 3B), implying that alpha lateralization had no direct causal role for producing the behavioral cueing effect. This result resembles that of Antonov et al. (2020) who found that the effect of endogenous spatial cues on alpha lateralization lagged behind the cueing effect on behavioral performance. Similarly, Bacigalupo and Luck (2019) found that alpha lateralization in a visual search task outlasted average response times and other electrophysiological indices of attention.

Moreover, there was no subsequent re-lateralization of alpha power (i.e. decrease ipsilateral/increase contralateral to cue) at latencies longer than 225 ms mirroring the behavioral IOR effect. This missing re-lateralization in the face of a strong behavioral inhibition effect is striking given the alleged role of alpha oscillations in attentional inhibition (Foxe and Snyder, 2011). Nonetheless, several other studies have also found no evidence for a direct role of alpha lateralization for inhibition of non-cued or irrelevant locations or stimuli (Noonan et al., 2016; Slagter et al., 2016; Antonov et al., 2020; Gundlach et al., 2020), suggesting that the suppressive account of alpha oscillations might have to be revisited (Slagter et al., 2016; Foster and Awh, 2019; van Moorselaar and Slagter, 2020). To conclude, the delayed emergence of alpha lateralization towards the cue and the missing subsequent re-lateralization mirroring the behavioral IOR suggest that cue-induced alpha-band lateralization may not be directly involved in producing behavioral effects of exogenous spatial orienting, or in modulating the neural response to target stimuli. Rather, alpha-lateralization might be associated with other processes coinciding with cueing, or with salient events in general, such as saccadic eye movements.

While subjects generally followed the instruction to maintain steady central fixation, they frequently made miniature eye movements (< 1 dva), and the direction of these microsaccades was consistently biased away from the cued location. This bias is probably a consequence of the inhibition of larger, reflexive saccades toward the exogenous cue (Rolfs et al., 2004). Notably, this compensatory microsaccade bias peaked at about the same time when alpha was maximally lateralized *towards* the cued location (Fig. 3A, D). Even though these effects were tuned in opposite directions, they were positively correlated: stronger alpha lateralization towards the cue was strongly associated with *reduced* microsaccade bias away from the cue.

The relationship between saccadic and microsaccadic eye movements and the alpha rhythm has long been established (Gaarder et al., 1966; Ulrich, 1990; Dimigen et al., 2009; Staudigl et al., 2017). Recently, interest in this relationship has been renewed based on studies showing that cue-induced alpha lateralization is stronger in trials with cue-driven microsaccades (Liu et al., 2022a), and that microsaccades are followed by alpha lateralization transients (Liu et al., 2022b). Importantly, Liu et al. (2022a) observed cue-induced alpha lateralization even in the absence of a microsaccade, suggesting that alpha lateralization is associated with both executed as well as sub-threshold, non-executed microsaccades, thus representing two aspects of oculomotor behavior (Liu et al., 2022a). In a similar vein, Popov et al. (2021) have demonstrated that alpha power modulations are strongly coupled with eye movements, and that the scalp topographies of saccade-related alpha power modulations are consistent with saccade direction. Given that eye movements are often biased towards cued or task-relevant locations, and given that miniature eye movements can easily escape fixation monitoring and EEG artifact removal, this may explain the high topographical specificity of alpha topographical distributions in tasks using spatial cueing (Rihs et al., 2007), visual search (Samaha et al., 2016; Foster et al., 2017), and visual working memory (Foster et al., 2016). Hence, we propose that cue-induced alpha lateralization in this study did not reflect a mechanism subserving the modulation of neuronal responses to target stimuli, but a function of the oculomotor system. Given that task instructions demanded steady fixations, pro-saccades had to be suppressed, as indicated by microsaccade bias in the opposite direction. The positive correlation between alpha lateralization and microsaccade bias (i.e. reduced bias away from the cue when alpha lateralization towards the cue was strong) suggests that both phenomena reflect the preparation for non-executed, reflexive pro-saccades towards the exogenously cued location.

This oculomotor interpretation is partially consistent with previous accounts of the role of alpha oscillations for cortical excitability and inhibition (Klimesch et al., 2007; Jensen and Mazaheri, 2010; Foxe and Snyder, 2011) by assuming that alpha-band lateralization is involved in controlling excitability in anticipation of saccade execution and landing. Given the tight coupling between eye movements and shifts of attention, this involvement would be consistent with the effect of spontaneous fluctuations of alpha-band lateralization and spatial attention (Bengson et al., 2014) or subjective contrast appearance (Balestrieri and Busch, 2022). Moreover, given that most cueing paradigms are designed such that attention is oriented towards the cued location, cue-induced alpha lateralization will typically show lateralization towards the attended location, even though it may not directly serve the neural amplification of attended targets or inhibition of unattended input. The IOR paradigm, in which attention is oriented away from the cued location at longer latencies makes it possible to reveal that alpha lateralizes towards the salient cue stimulus, not towards the direction of attentional (re)orienting.

## Conclusion

The current study shows that alpha lateralization does not track the time course and direction of attentional orienting in the inhibition of return (IOR) paradigm with exogenous cues. However, the time course of alpha lateralization towards the cue was very similar to that of microsaccades away from the cue.

This set of results, together with recent evidence from the literature, suggests that alpha lateralization and microsaccades are both consequences of planned, but ultimately suppressed, saccades toward the cued position. In this framework, alpha lateralization would modulate cortical excitability in anticipation of the saccade itself, whereas the microsaccades in the direction opposite to the cue would be reactively generated by suppressing the saccade toward the cue. This claim is further strengthened by a covariation observed between the two variables: the stronger the microsaccadic rate away from the cue, the weaker the alpha lateralization toward the cue.

## Acknowledgments

This work was supported by grants from the German Research Foundation (DFG; BU2400/8-1 and BU 2400/9-1) to NAB. We thank Teresa Berther, Johanna Seroka and Emma Brüssau for their help with data acquisition.

## Pregistration and Data availability

It is possible to access the pre-registration of the present study at the following link: https://aspredicted.org/blind.php?x=MWH_Y88.

Data and all the code for the present study will be made publicly available upon publication of the paper.

